# The rewarding properties of safety signals established by a two-way active avoidance task in male rats

**DOI:** 10.1101/2025.10.09.681387

**Authors:** Laura Vercammen, Tom Beckers, Bram Vervliet, Laura Luyten

## Abstract

Actively avoiding threatening stimuli or situations is an adaptive defensive strategy that minimizes the risk of potential harm. Despite the prominent role avoidance plays in our daily lives, the mechanisms that reinforce avoidance behavior remain incompletely understood. Accumulating evidence implicates the mesolimbic dopamine reward system in the acquisition of avoidance, suggesting that the omission of an aversive event may acquire rewarding properties and function as a positive reinforcer of avoidance behavior. In a series of six experiments (N = 246 male Wistar rats), we examined whether safety signals that coincide with successful avoidance responses in a two-way active avoidance (2WAA) task facilitate avoidance acquisition and acquire reward-like properties, by assessing whether they elicit approach behavior and support the acquisition of a novel instrumental response. Although the presentation of a safety signal accelerated avoidance learning in some experiments, the effect was not consistent. Furthermore, the safety signal neither elicited approach behavior nor supported the acquisition of a novel instrumental response. Together, these findings question whether safety signals in avoidance learning acquire rewarding properties detectable through behavioral measures.

## 1. Introduction

Actively avoiding threatening stimuli or situations serves as an adaptive behavioral strategy for reducing potential danger in our everyday life (Krypotos et al., 2015). For example, we typically refrain from walking in the path of oncoming traffic to minimize the risk of physical harm, or will take an alternative route to work to avoid encounters with unleashed dogs in the park. While avoidance is a prominent part of our everyday behavior, there is ongoing debate about the mechanisms that reinforce it (Fisher & Urcelay, 2024). Avoidance is reinforced in the absence of an explicit observable event, as the performance of an avoidance action results in the omission of a potential threat (Cain, 2019). Various theoretical models have been developed that differ in how they interpret the role of the aversive event’s absence in reinforcing avoidance behavior. Negative reinforcement theories (e.g., Mowrer’s two-factor theory, Mowrer & Lamoreaux, 1946) suggest that avoidance is reinforced by the reduction in fear that accompanies the avoidance action. This reduction in fear associated with the avoidance response will ensure that this action is repeated in the future. In contrast, alternative perspectives conceptualize avoidance as being positively reinforced. The omission of the US is perceived as a rewarding or “better-than-expected” outcome, as safety is obtained instead of the expected threat, thereby positively reinforcing the avoidance behavior (Fernando et al., 2014; Cain, 2019; Dinsmoor, 2001).

Experimental models, both in humans (e.g., Krypotos et al., 2018) and rodents (e.g., Servatius et al., 2008; Bolles et al., 1966) have been developed to investigate how the omission of the aversive event can reinforce avoidance behavior. In rodent research, the two-way active avoidance (2WAA) task is a widely used paradigm for studying the neural mechanisms underlying avoidance acquisition (Fernández-Teruel & Tobeña, 2020). In this task, a conditioned stimulus (CS, e.g., a tone) is repeatedly paired with an aversive unconditioned stimulus (US, e.g., a foot shock). Rats can avoid the presentation of the US by shuttling to the opposite compartment of a shuttle box during the presentation of the CS. While preclinical models have identified important brain areas involved in the acquisition of avoidance (Choi et al., 2010; Moscarello & Ledoux, 2013; Ramirez et al., 2015; Martinez et al., 2013), there is still ongoing debate about the mechanisms that reinforce it (Fisher & Urcelay, 2024). Is avoidance negatively reinforced because it reduces the fear that is elicited by the CS, or is it positively reinforced by the rewarding nature of the safety it produces?

Accumulating evidence supports the positive reinforcement framework, because of the involvement of the mesolimbic dopamine reward system in avoidance learning, which plays a critical role in reward processing and appetitive reinforcement learning (Schultz, 2016). Pharmacological blockade of dopamine receptors, both systemically or in the nucleus accumbens (NAc), an important locus within the reward system (Day & Carelli, 2007), prior to avoidance learning impairs its acquisition (Wietzikoski et al., 2012; Reis et al., 2004; Boschen et al., 2011). Moreover, restoring dopamine signaling in the amygdala and striatum of dopamine-deficient mice is necessary for successfully acquiring the avoidance response (Darvas et al., 2011). Further converging evidence comes from a study that directly measured dopamine release in the NAc during a lever-press avoidance task. Specifically, increases in DA were observed during safety signals that signaled the exact moment that the aversive US was omitted or terminated, a pattern similar to the DA response typically elicited by stimuli predicting reward (Oleson et al., 2012; Schultz, 2016), suggesting that the omission of the US may elicit a similar burst in dopamine as a rewarding stimulus. Importantly, dopamine release does not automatically indicate that a stimulus is rewarding. While dopamine is mostly associated with reward processing, it does respond do a broader range of salient experiences, including non-rewarding and aversive events (Bromberg-Martin et al., 2010; Baik, 2020), and therefore a central role of dopamine is consistent with punishment-based models of avoidance as well as (Amin et al., 2025). Hence, the aim of the current study was to evaluate whether stimuli that signal the successful omission or termination of the aversive US (i.e., so-called safety signals) can trigger reward-like behavior as observed for stimuli that predict the receipt of natural rewards (Berridge & Robinson, 2003). If safety signals elicit behavioral responses similar to those observed for appetitive cues, this would support the interpretation that the observed dopamine activity reflects a reward process.

There is some initial evidence that suggests that stimuli that signal omissions of aversive USs are able to trigger reward-like behavior. When rodents passively experience omissions of an aversive foot shock in a conditioned inhibition procedure, safety signals related to these omissions are able to trigger approach behavior in mice (Rogan et al., 2005), and promote food seeking behavior in rats (Walasek et al., 1995). Although this would support the notion that safety signals acquire rewarding properties, Fernando et al. (2013) found that these signals were unable to promote the acquisition of a novel instrumental response, in contrast to what has been observed for appetitive cues (Fernando et al., 2013). This failure to observe conditioned reinforcing effects of the safety signal might have been caused by the lack of control experienced by the subjects when safety signals are established via conditioned inhibition. Active avoidance procedures on the other hand, enable subjects to experience active control over US omission, which might enhance the reinforcing effects of safety signals. Indeed, in a follow-up study, Fernando et al. (2014) observed that safety signals acquire reinforcing properties when using an unsignaled lever-press avoidance task. Rats preferentially responded on a lever that avoided the US and produced a safety signal, compared to a lever that did not produce a safety signal while also omitting the US. Moreover, it has been repeatedly reported that safety signals can accelerate avoidance learning, again hinting towards a positive reinforcement mechanism (e.g., jump-up avoidance, lever-press avoidance; Cándido et al., 1991; Dillow et al., 1972). But, while safety signals acquire reinforcing properties, it is not yet clear whether they elicit similar behavioral responses as appetitive cues that signal reward.

In a series of six experiments, we evaluated whether safety signals acquire reinforcing and rewarding properties. Specifically, we evaluated whether safety signals accelerate two-way active avoidance learning, as has previously been observed in other avoidance tasks (Cándido et al., 1991, Dillow et al., 1972), and whether they trigger reward-like behavior. We hypothesized that safety signals, through their association with the omission of the US, gain rewarding properties and thereby elicit similar behavioral responses as observed for cues that signal natural rewards. Following three to five days of avoidance training, which did or did not include the safety signal, rats were subjected to a compartment-preference (CP) test (modified from Rogan et al., 2005) and/or a lever-preference (LP) test (modified from Fernando et al., 2013) to measure whether a safety signal elicits approach behavior and whether it can promote the acquisition of a novel instrumental response, respectively.

## 2. Methods and Materials

### 2.1. Preregistration and data availability

All experiments were preregistered on the Open Science Framework (https://doi.org/10.17605/OSF.IO/M7XJR). The raw data files, processed data files and R scripts generated for the current study are available via the same link. Video files are available upon request.

### 2.2. Subjects

A total of 246 male Wistar rats (weighing between 275 – 300 g upon arrival in the lab) was purchased from Janvier Labs (Le Genest Saint Isle, France). Rats were housed per 2-4 in standard animal cages with cage enrichment and kept under conventional laboratory conditions (12-h day-night cycle; lights on 7:00-19:00; 22°C) with ad libitum access to food and water. All samples sizes were based on power analyses, calculated using G*Power 3.1.9.7 (power = .80, alpha = .05) and effect sizes available in the literature. If no prior effect sizes were available, an intermediate effect size was chosen based on a systematic review paper that calculated the mean effect size of 410 experiments in the field of rodent fear learning (Carneiro et al., 2018). Experiments were conducted between 8:00 and 18:00. Rats were handled briefly on two consecutive days, prior to behavioral testing. All experiments were approved by the KU Leuven animal ethics committee (project P141/2021) and conducted in accordance with the Belgian and European laws, guidelines and policies for animal experimentation, housing and care (Belgian Royal Decree of 29 May 2013 and European Directive 2010/63/EU on the protection of animals used for scientific purposes of 20 October 2010).

### 2.3. Apparatus

#### Two-way active avoidance (Coulbourn Instruments setup)

For experiments 1, 2, 3, 6 and supplemental experiment A (see Supplementary Materials), two-way active avoidance training took place in two identical rectangular shuttle boxes (50.8 cm x 25.4 cm x 30.5 cm), each placed inside a larger sound-attenuating cubicle (72.4 cm x 41.1 cm x 43.2 cm, Coulbourn Instruments, Pennsylvania, USA). The side walls and ceilings of the shuttle boxes were made of metal, whereas the front and back walls were clear Plexiglas. The shuttle box was divided into two equal compartments by a metal divider placed halfway along the length of the shuttle box. Rats could freely move from one side of the box to the other via a passage (8 cm width) in the divider. The floor of the shuttle box consisted of a stainless steel grid, through which scrambled foot shocks could be delivered (0.6 mA, max 10 s) that served as the unconditioned stimulus (US). In each compartment of the shuttle box, a speaker was mounted to the side wall to deliver a 3-kHz tone (presented at +/- 75 dB) that served as the conditioned stimulus (CS). A 0.5-W white house light was mounted to the ceiling of each compartment and served as a safety signal. When the safety signal was not on, the shuttle box was unilluminated. Two photocell sensor bars, one in each side of the shuttle box, ensured automatic detection of the position of the rat throughout the experiment. The delivery of the stimuli and data collection was fully automated by a PC running Graphic State 4 software (Coulbourn Instruments). All behavioral sessions were videorecorded via an infrared camera (HD IP camera, Foscam C1, Shenzhen, China) mounted to the ceiling of the isolation cubicle.

#### Two-way active avoidance (Med Associates setup)

For experiments 4 and 5, two-way active avoidance training took place in four identical rectangular shuttle boxes (57.4 cm x 28.7 cm x 24.13 cm), each housed inside a larger sound-attenuating cubicle (96.5 cm x 40.6 cm x 55.9 cm, Med Associates, Vermont, USA). The side walls of the shuttle boxes were made of metal, whereas the front and back walls, as well as the ceiling of the shuttles boxes were clear Plexiglas. The shuttle box was divided into two equal compartments by a metal divider placed halfway along the length of the shuttle box (Med Associates). Rats could freely move from one side of the box to the other via a passage (8 cm width) in the divider. The floor of the shuttle box consisted of a stainless steel grid, through which scrambled foot shocks could be delivered (0.6 mA, max 10 s) that served as the US. In each compartment of the shuttle box, a speaker was mounted to the side wall to deliver a 3-kHz tone (presented at +/- 75 dB) that served as the CS. A white protruding house light (55 lx) was mounted at the top of the side wall of each compartment, and a flat round yellow LED light (40 lx) was mounted at the bottom of the side wall of each compartment. The light cues were used as either the safety signal or the control cue (counterbalanced between rats). When the safety signal and control cue were not on, the shuttle box was unilluminated. Two photobeam sensor bars, one in each side of the shuttle box, ensured automatic detection of the position of the rat throughout the experiment. The delivery of the stimuli and data collection was fully automated by a PC running Med PC 5 (Med Associates). All behavioral sessions were videorecorded via an infrared camera (HD IP camera, Foscam C1, Shenzhen, China) mounted to the ceiling of the isolation cubicle.

#### Compartment-preference (CP) test

For experiments 2, 3 and supplemental experiment A, the compartment-preference (CP) tests took place in the same shuttle boxes as used for avoidance training, which were modified for the CP tests with regard to their tactile, visual and olfactory features (indicated in the figures with a grey background). Specifically, the front and back walls of the shuttle boxes were covered with an opaque black film and a red solid plastic floor was inserted to cover the electric grid. The context was scented with a pine-odored cleaning product (Bref Pine, 5.55%), by spraying 3 x in the drop pan. For experiments 4, 5 and 6, the CP tests also took place in the same shuttle boxes as used for avoidance training, without contextual modifications. Time spent in each compartment was automatically recorded through Graphic State 4 software (Coulbourn Instruments) or Med-PC 5 software (Med Associates).

#### Lever-preference (LP) test

The lever-preference (LP) test took place in the same shuttle boxes as used for avoidance training, which were modified for LP by inserting two levers (Med Associates) in the right side wall of the shuttle box. The number of lever presses on each lever was automatically recorded through Med-PC 5 software.

### 2.4. Behavioral procedures

Table 1 provides an overview of the behavioral sessions of each experiment. Supplemental Table 1 and 2 provide a detailed overview of the parameters used during the behavioral protocols. Supplemental Table 1 provides an overview of the parameters that were used during avoidance training, whereas Supplemental Table 2 provides an overview of the parameters that were used during the CP tests.

**Table 1.**
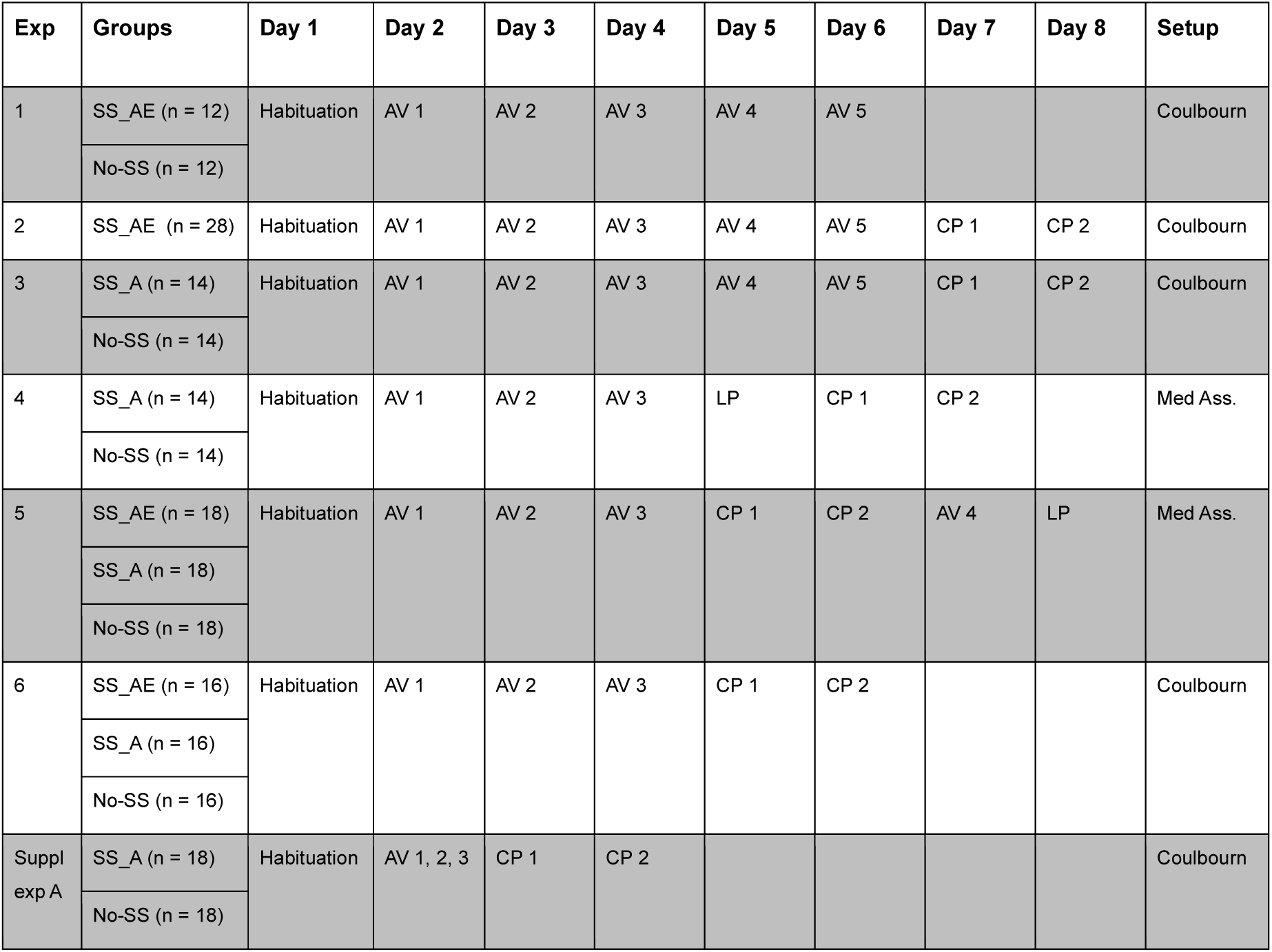
Overview of the experimental procedures. AV = avoidance training session, CP = compartment-preference test. LP = lever-preference test. Supplemental Experiment A can be found in the Supplemental Materials. SS_AE groups received the safety signal contingent upon avoidance and escape responses, SS_A groups received the safety signal only after successful avoidance responses, and no-SS groups did not receive the safety signal during avoidance learning.

#### 2.4.1. Experiment 1

##### Two-way active avoidance training

Upon arrival in the lab, rats were randomly assigned to either the SS_AE group (n = 12) or the no-SS group (n = 12). Twenty-four hours before the start of avoidance training, all rats were habituated to the shuttle box (Coulbourn instruments, Pennsylvania, USA) for 1 hour. Starting the following day, the rats were subjected to five consecutive daily avoidance training sessions (Vercammen et al., 2025). Each avoidance training session started with a 5-min acclimation period during which no stimuli were delivered. After acclimation, the rats were given 30 presentations of the CS and US, separated by an intertrial interval (ITI) averaging around 2 minutes (60s – 180s). The CS lasted a maximum of 20 seconds and was followed immediately by the US which lasted a maximum of 10 seconds. If the rat shuttled to the other compartment during CS presentation, the CS was terminated and no shock was given, i.e., the US was avoided and the trial was labeled as ending in an avoidance response. If the rat crossed to the other compartment during the presentation of the US, the shock was terminated and the trial was labeled as ending in an escape response. Upon each successful avoidance or escape response, rats of the SS_AE group received a safety signal that was presented in the compartment where the rat avoided or escaped into (house light on for 5 seconds). Rats of the no-SS group did not receive the safety signal. The total duration of each training session was 55 minutes. The number of avoidance responses, escape responses and escape failures (i.e., the rat remained in the compartment throughout the US presentation) were recorded via Graphic State 4 software (Coulbourn Instruments).

#### 2.4.2. Experiment 2

##### Two-way active avoidance training

Two-way active avoidance training proceeded as described above (Experiment 1), except that the ITIs were shortened to 60 seconds on average (30 s – 90 s), with the intention to make the task more difficult and augment the effect of the safety signal on avoidance learning (Song et al., 2017). All subjects (N = 28) underwent the avoidance protocol that included the safety signal after successful avoidance and escape responses (SS_AE).

##### Compartment-preference test 1

Twenty-four hours after the final avoidance training session, the rats were placed in the modified shuttle box for a 10-min session. The CP test started with a 5-min acclimation period during which no stimuli were presented. After five minutes, the safety signal was turned on in the compartment where the rat was at that point in time (from thereon referred to as the SS-compartment). For the following five minutes, whenever the rat left the SS-compartment, the safety signal was turned off. Upon re-entering the SS-compartment, the safety signal was turned back on. Time spent in each compartment was automatically recorded.

##### Compartment-preference test 2

Twenty-four hours after CP test 1, the rats were again placed in the modified shuttle box for a 5-min session. After 5 seconds, the safety signal turned on in the SS-compartment (same compartment as during CP test 1). Similar as in CP test 1, whenever the rat was in the SS-compartment, the SS was on. Time spent in each compartment was automatically recorded.

#### 2.4.3. Experiment 3

##### Two-way active avoidance training

Upon arrival in the lab, rats were randomly assigned to either the SS_A group (n = 14) or the no-SS group (n = 14). Two-way active avoidance training proceeded as described above (Experiment 1), except that the ITIs were shortened to 60 seconds on average (30 s – 90 s), as in Experiment 2. Moreover, rats of the SS_A group received the safety signal after each avoidance response, but not after escape responses. This change was introduced to ascertain that safety signals acquire rewarding properties via omission of the US (i.e., during avoidance responses), and not by termination of the US (i.e., during escape responses), which has also been found to activate reward-related systems (Navratilova et al., 2015). Rats of the no-SS group did not receive the safety signal during avoidance training.

##### Compartment-preference test 1

Twenty-four hours after the final avoidance training session, the first CP test took place as described above (Experiment 2), except that the SS-compartment assignment was no longer dependent on the behavior of the rat. Instead, the SS-compartment was predetermined and counterbalanced between rats. This randomization was implemented to control for the potentially biased safety signal-compartment assignment when the safety signal is turned on dependent on the behavior of the rats.

##### Compartment-preference test 2

Twenty-four hours later, CP test 2 took place as described above (Experiment 2), except that the safety signal turned on in the opposite compartment as in CP test 1 (i.e., if the left compartment was the SS-compartment during CP test 1, the right compartment was the SS-compartment during CP test 2). This change was implemented to ensure that safety signal preference is measured, instead of preference for a specific compartment. If the safety signal elicits approach behavior, rats should follow the safety signal, independent of the compartment in which it is presented.

#### 2.4.4. Experiment 4

##### Two-way active avoidance training

For this experiment, a Med Associates setup was used instead of the Coulbourn setup, for purely practical reasons, i.e., the availability of four shuttle boxes instead of two. Upon arrival in the lab, rats were randomly assigned to either the SS_A group (n = 14) or the no-SS group (n = 14). Twenty-four hours before the start of avoidance training, all rats were habituated to the shuttle box (Med Associates, Vermont, USA) for 1 hour. On the following day, the rats were subjected to three consecutive daily avoidance training sessions. The number of training sessions was reduced from 5 to 3 in comparison with the preceding experiments because rats reached asymptotic performance by the third training session in the previous experiments. Each avoidance training session started with a 10-min acclimation period, during which the rats were habituated to a control cue (white protruding house light or flat yellow light, counterbalanced between rats). The control cue was presented 5 times for 10 seconds (average ITI ∼65 s) to ensure that both groups habituated to the control light. Moreover, the no-SS group also received 5 presentations of the safety signal light to ensure that they were also habituated to this cue (average ITI ∼25 s), and thereby eliminate potential novelty effects during the CP tests. The presentation schedule of the cues can be found in the Supplementary Materials (Suppl. Table 3). After the 10-min acclimation, the experiment proceeded as described for Experiment 3, except that the safety signal given after each avoidance response in the SS_A group, was presented in both compartments simultaneously and was prolonged to 10 seconds (instead of 5) to attract maximum attention to the signal. The total duration of the training session was 45 minutes.

##### Lever-preference test

Twenty-four hours after the final avoidance training session, rats were reintroduced for a 30-min session to the shuttle box that now contained two levers mounted on the right side wall. Pressing on one of the levers produced the safety signal for 5 seconds, whereas pressing on the other lever produced the control cue for 5 seconds (counterbalanced between the left and right lever). The number of lever presses on each lever was recorded.

##### Compartment-preference test 1

Twenty-four hours after the LP test, the rats were reintroduced to the shuttle box for a 5-min session. In contrast to previous experiments, the original shuttle box without contextual modifications was used, with the intention of enhancing the safety signaling properties of the safety signal, by placing them in the original threat context (Hefner et al., 2017). After 5 seconds, the safety signal turned on in one of the compartments (SS-compartment), and the control cue in the other compartment (CC-compartment; counterbalanced between rats). After the first shuttle, the cue in the compartment that the rat left was turned off, so that only the cue in the compartment where the rat shuttled into remained on. From there onwards, whenever the rat was in the SS-compartment, the safety signal was turned on and the control cue off, and when the rat was in the CC-compartment, the control cue was turned on and the safety signal off. Time spent in each compartment was recorded, starting from the first shuttle onwards.

##### Compartment-preference test 2

Twenty-four hours after CP test 1, the rats were reintroduced into the shuttle box for a 5-min session. The second CP test proceeded similar to CP test 1, except that the SS- and CC-compartments were switched (e.g., if the left compartment was the SS-compartment during CP test 1, the right compartment was the SS-compartment during CP test 2).

#### 2.4.5. Experiment 5

##### Two-way active avoidance training

Upon arrival in the lab, rats were randomly assigned to either the SS_A group (n = 18), the SS_AE group (n = 18) or the no-SS group (n = 18). The SS_AE group (which receives the safety signal after avoidance and escape responses, and which was already used in Experiments 1 and 2) was added to the design, to characterize the relative importance of avoidance vs. the combination of avoidance and escape responses in assigning rewarding properties to the safety signal. Two-way active avoidance training further proceeded as in Experiment 4.

##### Compartment-preference test 1

Twenty-four hours after the final avoidance training session, the rats were reintroduced into the shuttle box for a 10-min session. We again slightly changed the procedure of the CP test in order to have a clear and unbiased read-out. As in Experiment 4, one of the compartments was assigned as the SS-compartment, and the other compartment as the CC-compartment (counterbalanced between rats). In contrast with Experiment 4, the session started with a 5-min acclimation period during which the rats were allowed to freely move between the compartments. After 5 minutes, the relevant cue was turned on in the compartment where the rat was at that point in time (e.g., if the rat was in the left compartment after 5 minutes and this was the SS-compartment, the SS was turned on). From there onwards, whenever the rat left the SS-compartment, the safety signal was turned off and the control cue was turned on in the other compartment. Time spent in each compartment was recorded, starting from the first shuttle onwards.

##### Compartment-preference test 2

Twenty-four hours after CP test 1, the rats were reintroduced into the shuttle box for a 5-min session. The second CP test proceeded similar to CP test 1, except that the SS- and CC-compartments were switched (e.g., if the left compartment was the SS-compartment during CP test 1, the right compartment was the SS-compartment during CP test 2).

##### Two-way active avoidance training

Twenty-four hours after CP test 2, all rats were given a single avoidance training session, which proceeded as described above for this experiment. This training session was added to prevent extinction of the safety signaling properties of the safety signal.

##### Lever-preference test

Twenty-four hours after the fourth avoidance training session, rats were subjected to a lever-preference test, as described in Experiment 4.

#### 2.4.6. Experiment 6

##### Two-way active avoidance training

Upon arrival in the lab, rats were randomly assigned to either the SS_A group (n = 16), the SS_AE group (n = 16) or the no-SS group (n = 16). All procedures were equal to those of Experiment 5, except that the experiment took place in the Coulbourn Instruments setup, instead of the Med Associates setup, to evaluate potential setup-dependent effects.

##### Compartment-preference test 1

Twenty-four hours after the final avoidance training session, the rats were subjected to CP test 1. This test proceeded similar as CP test 1 in Experiment 5.

##### Compartment-preference test 2

Twenty-four hours after CP test 1, the rats were subjected to CP test 2. This test proceeded similar as CP test 2 in Experiment 5.

### 2.5. Statistical analyses

All analyses were preregistered on the Open Science Framework, unless stated otherwise. Data are expressed as means ± SEM and analyzed using R statistical software (version 3.5.1). The mean number of avoidance responses was calculated for each session per group and analyzed using mixed design ANOVAs (repeated-measures factor: Session, between-subjects factor: Group). Moreover, the mean number of avoidance responses was calculated per block of 10 trials within the first avoidance training session and analyzed using a mixed design ANOVA (repeated-measures factor: Block, between-subjects factor: Group). For the compartment-preference test, time spent in the SS-compartment was evaluated using one-sample t-tests for rats of the safety signal groups (SS_AE and SS_A, > 50 %) and the no-SS group (≠ 50%). Moreover, independent samples t-tests were used to evaluate whether time spent in the SS-compartment was significantly elevated for rats of the safety signal groups, compared to the no-SS group (SS_AE, SS_A > no-SS). For the lever-preference test, a percentage SS preference (number of SS lever presses / total number of lever presses * 100) was calculated and analyzed using one-sample t-tests for rats of the safety signal groups (> 50%) and rats of the no-SS group (≠ 50%). Preregistered secondary analyses can be found in the Supplementary Materials.

## 3. Results

### 3.1. Experiment 1

The aim of Experiment 1 was to evaluate whether 2WAA acquisition is accelerated when a safety signal follows each successful avoidance or escape response (Cándido et al., 1991). Rats were assigned to either the SS_AE group (n = 12), where a safety signal (house light for 5 s) followed each avoidance and escape response, or to the no-SS group (n = 12), where a safety signal was not presented.

The number of avoidance responses (ARs) significantly increased with repeated training, as illustrated by a significant effect of Session (*F*(2.55,56.19) = 44.856, *p* < .001, ƞ*_p_²* = 0.671; Figure 1A). When taking into account all five training days, there was no significant main effect of Group (*F*(1,22) = 2.292, *p* = .144, ƞ*_p_²* = 0.094), nor a significant Group by Session interaction effect (*F*(2.55,56.19) = 0.903, *p* = .432, ƞ*_p_²* = 0.039), suggesting that, overall, avoidance learning occurred similarly for both groups (Figure 1A). When exploratively analyzing the data of the first avoidance training session in three blocks of 10 trials each, we observed a significant main effect of Block (*F*(2,44) = 4.678, *p* = .014, ƞ*_p_²* = 0.175), but no main effect of Group (*F*(1,22) = 2.245, *p* = .148, ƞ*_p_²* = 0.093), nor a significant Group by Block interaction effect (*F*(2,44) = 0.351, *p* = .706, ƞ*_p_²* = 0.016, Figure 1B). Together, these findings indicate that the safety signal did not significantly enhance avoidance learning.

**Figure 1.**
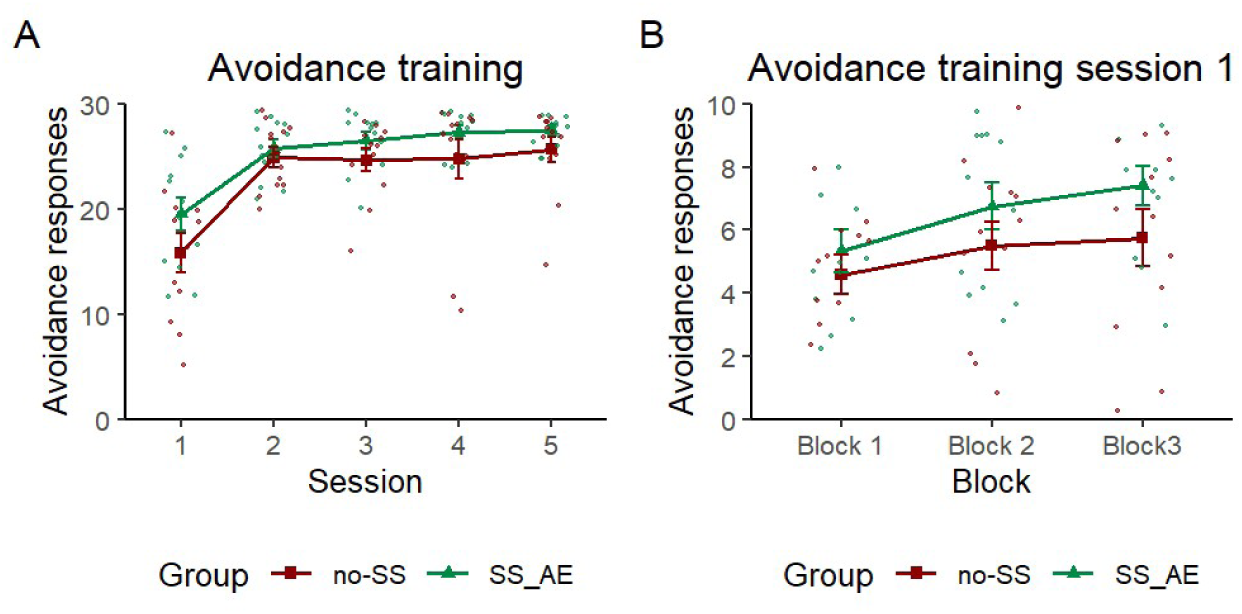
Experiment 1 avoidance training. (A) Mean (± SEM) number of ARs for the SS_AE (n = 12) and no-SS (n = 12) groups. (B) Mean (± SEM) number of ARs per group during the first avoidance training session in 3 blocks of 10 trials each.

### 3.2. Experiment 2

The aim of Experiment 2 was to establish a behavioral protocol to evaluate whether safety signals acquire rewarding properties after avoidance learning. All subjects (N = 28) underwent five days of 2WAA training which included the safety signal, followed by a compartment-preference (CP) test to evaluate whether the safety signal triggered approach behavior.

The number of ARs significantly increased with repeated training, as illustrated by a significant effect of Session (*F*(1.68,45.44) = 46.741, *p* < .001, ƞ*_p_²* = 0.634; Figure 2A). During the first five minutes of habituation of CP test 1, rats showed no significant preference for a particular side of the box (*t*(27) = 1.957, *p* = .061, *d* = 0.370), although there was a trend towards preference for the right compartment of the contextually modified shuttle box. Once the safety signal was introduced, rats preferred the compartment where the safety signal was presented over the compartment where no stimulus was presented (*t*(27) = 2.890, *p* = .004, *d* = 0.546; Figure 2B). Similarly, during CP test 2, rats preferred the compartment where the safety signal was presented, compared to the no-stimulus compartment (*t*(27) = 2.900, *p* = .004, *d* = 0.548; Figure 2C). Together, these findings suggest that the safety signal may elicit approach behavior, as previously observed for cues that predict natural rewards (Berridge & Robinson, 2003).

**Figure 2.**
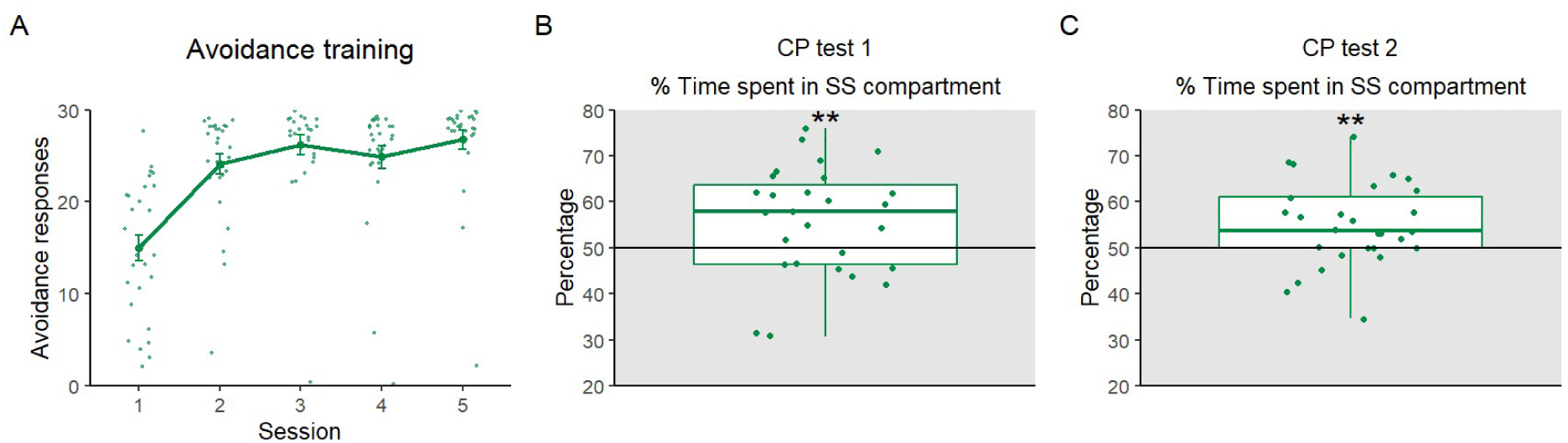
Experiment 2 avoidance training and compartment-preference (CP) tests. (A) Mean (± SEM) number of ARs. Boxplot of the percentage of time spent (B) in the SS-compartment of CP test 1, and (C) in the SS-compartment of CP test 2. ** *p* < .01.

### 3.3. Experiment 3

Experiment 3 was set up to address the limitations of Experiment 2 (lack of a no-SS control group and potentially biased SS-compartment assignment of CP tests). Moreover, in Experiments 1 and 2, the safety signal was presented after both avoidance and escape responses. Given our primary interest in the reinforcing role of US omissions (i.e., as observed during an avoidance response), we modified our approach and presented the safety signal only after avoidance responses in Experiment 3. Rats were randomly assigned to the SS_A group (n = 14), which received the safety signal after avoidance responses, or the no-SS group (n = 14), which did not receive the safety signal upon avoidance. Following five days of avoidance training, both groups were subjected to the CP tests.

One subject from the SS_A group was post-hoc excluded from the experiment due to a high number of escape failures in the first training session (26 failures out of 30 trials). We observed a significant main effect of Session (*F*(2.59,64.85) = 27.630, *p* < .001, ƞ*_p_²* = .525; Figure 3A), suggesting that the number of ARs increased with repeated training. The main effect of Group failed to reach significance (*F*(1,25) = 3.906, *p* = .059, ƞ*_p_²* = .135), despite a numerical trend for rats in the SS_A group to exhibit more avoidance responses, compared to rats in the no-SS group. When restricting the analysis to the block data of the first avoidance training session, the main effect of Group was significant (*F*(1,25) = 6.063, *p* = .021, ƞ*_p_²* = .195; Figure 3B). Post-hoc testing showed that rats from the SS_A group showed, on average, more avoidance responses in the first training session, compared to rats of the no-SS group (*t* = 2.462, *p* = .021), in line with the hypothesis that a safety signal accelerates avoidance learning.

**Figure 3.**
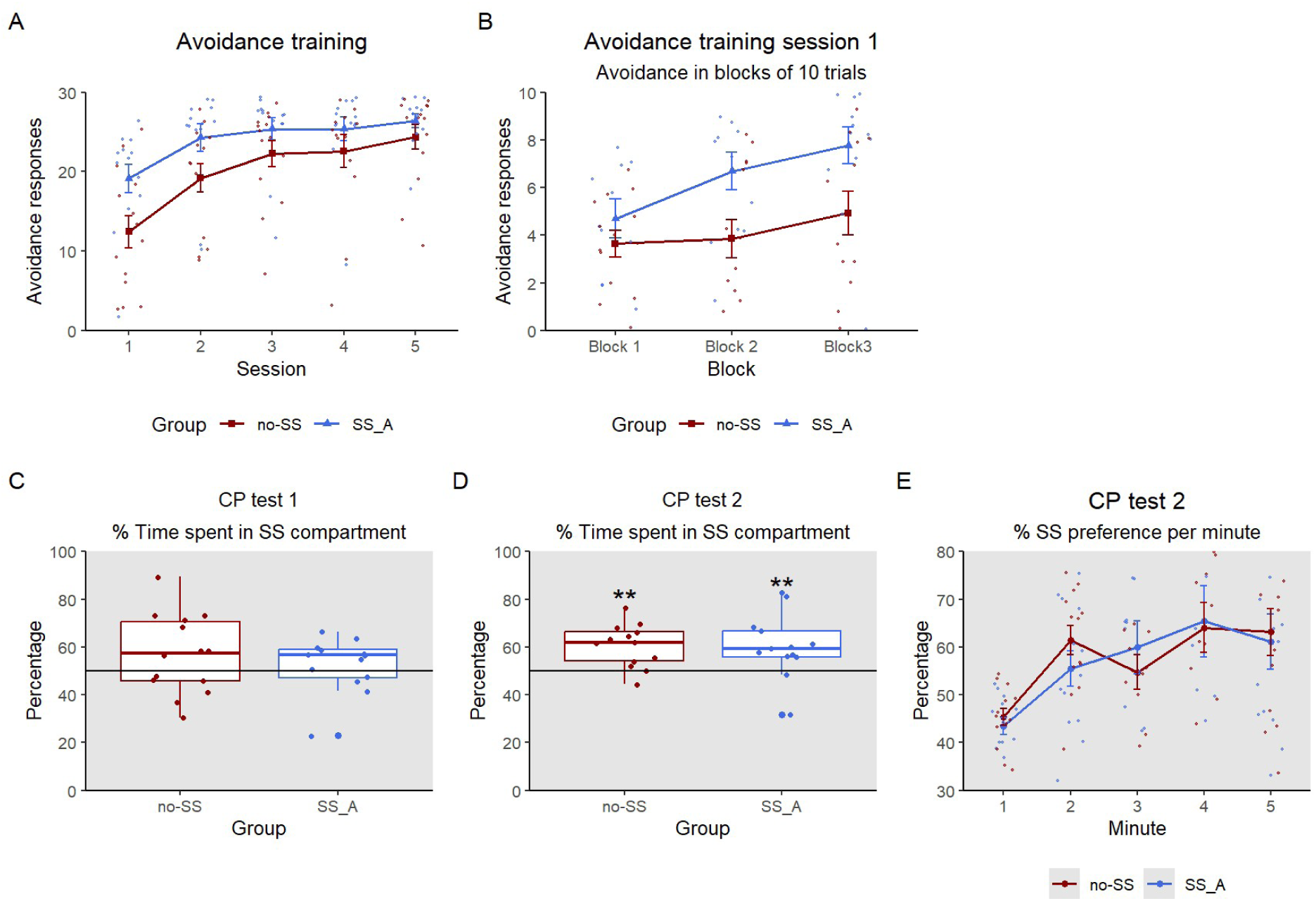
Experiment 3 avoidance training and compartment-preference tests. (A) Mean (± SEM) number of ARs for the SS_A (n = 14) and no-SS (n = 14) groups. (B) Mean (± SEM) number of ARs of the first avoidance training session in blocks of 10 trials for both groups. (C) Mean percentage time spent in the SS-compartment of CP test 1 for rats of both groups. (D) Mean percentage time spent in the SS-compartment of CP test 2 for both groups. ** *p* < .01. (E) Mean percentage time spent in the SS-compartment per minute of CP test 2 for both groups.

During the first 5 minutes of CP test 1 (habituation), rats showed no significant preference for a particular side of the box (SS_A group: *t*(12) = - 1.649, *p* = .125, *d* = −0.457; no-SS group: *t*(13) = 1.401, *p* = .185, *d* = 0.374). Once the SS was introduced, rats of both groups did not prefer the compartment where the SS was presented over the compartment where no stimulus was presented (SS_A group: *t*(12) = 0.779, *p* = .226, *d* = 0.216; no-SS group: *t*(13) = 1.536, *p* = .148, *d* = 0.411; Figure 3C), nor was there a significant difference between both groups in time spent in the SS-compartment (*t*(25) = −0.796, *p* = .783, *d* = −0.307). During CP test 2, both groups did spend significantly more time in the SS-compartment compared to the no-stimulus compartment (SS_A group: *t*(12) = 2.858, *p* = .007, *d* = 0.793; no-SS group: *t*(12) = 4.249, *p* = .001, *d* = 1.178; Figure 3D). When exploratively analyzing the % time spent in the SS-compartment during CP test 2 per minute, there was a significant main effect of Minute, suggesting that time spent in the SS-compartment increased throughout the test (*F*(4,96) = 7.863, *p* < .001, ƞ*_p_²* = .247, Figure 3E), and this increase was similar for both groups, as evidenced by a nonsignificant main effect of Group (*F*(1,24) = 0.025, *p* = .876, ƞ*_p_²* = 0.001) and a nonsignificant Group by Minute interaction (*F*(4,96) = 0.555, *p* = .696, ƞ*_p_²* = 0.023). These findings question whether the SS triggered approach behavior for rats of the SS_A group, given the absence of a significant SS-compartment preference during CP test 1 and the lack of a group difference for the SS-compartment preference that was observed during CP test 2. The increasing approach of the no-SS rats towards the safety signal during CP test 2 might reflect novelty-seeking, as they did not yet experience the safety signal before the start of the CP tests.

### 3.4. Experiment 4

The aim of Experiment 4 was to address the limitations of Experiment 3 by habituating the no-SS group to the safety signal during avoidance training to remove any novelty-induced effects during the CP tests. Moreover, a control cue was included in the experimental design to ascertain that the effects of the CP tests would not be driven by preference for a “light” (i.e., safety signal compartment) over a “dark” (i.e., no stimulus compartment) environment. Rats were randomly assigned to the SS_A group (n = 14) or the no-SS group (n = 14) upon arrival in the lab.

We observed a significant main effect of Session, suggesting that the number of ARs increased with repeated training (*F*(2,52) = 43.216, *p* < .001, ƞ*_p_²* = 0.624), but no significant main effect of Group (*F*(1,26) = 1.729, *p* = .200, ƞ*_p_²* = 0.062), nor a significant Group by Session interaction effect (*F*(2,52) = 2.597, *p* = .084, ƞ*_p_²* = 0.091; Figure 4A). Visual inspection of Figure 5A does suggest that SS_A rats may have already reached the learning asymptote during Session 2, in contrast with the no-SS rats reaching it in Session 3, but this was not statistically corroborated. When restricting the analysis to the data of the first avoidance training session, similarly, there was only a significant effect of Block (*F*(2,52) = 24.193, *p* < .001, ƞ*_p_²* = 0.482), but no significant effect of Group (*F*(1,26) = 0.001, *p* = .975, ƞ*_p_²* < .001), nor a significant Group by Block interaction effect (*F*(2,52) = 2.332, *p* = .107, ƞ*_p_²* = 0.082; Figure 4B).

**Figure 4.**
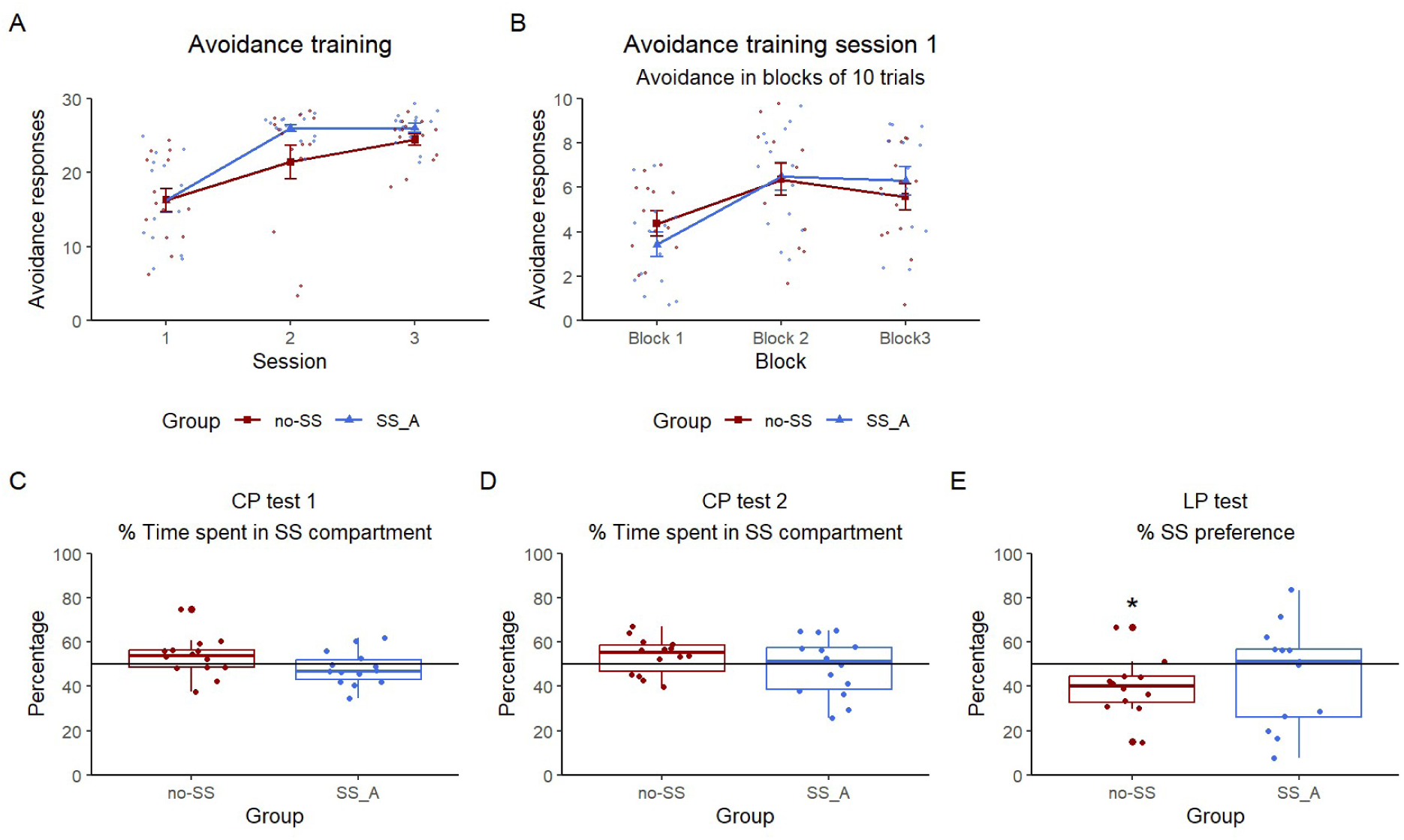
Experiment 4 avoidance training, compartment-preference and lever-preference tests. (A) Mean (± SEM) number of ARs for the SS_A (n = 14) and no-SS (n = 14) groups. (B) Mean (± SEM) number of ARs of the first avoidance training session in blocks of 10 trials for both groups. (C) Boxplot of the percentage time spent in the SS-compartment of CP test 1 for rats of both groups. (D) Boxplot of the percentage time spent in the SS-compartment of CP test 2 for both groups. (E) Boxplot of the percentage SS preference of the LP test for both groups. Rats of the no-SS group significantly preferred the CC lever. * *p* < .05.

**Figure 5.**
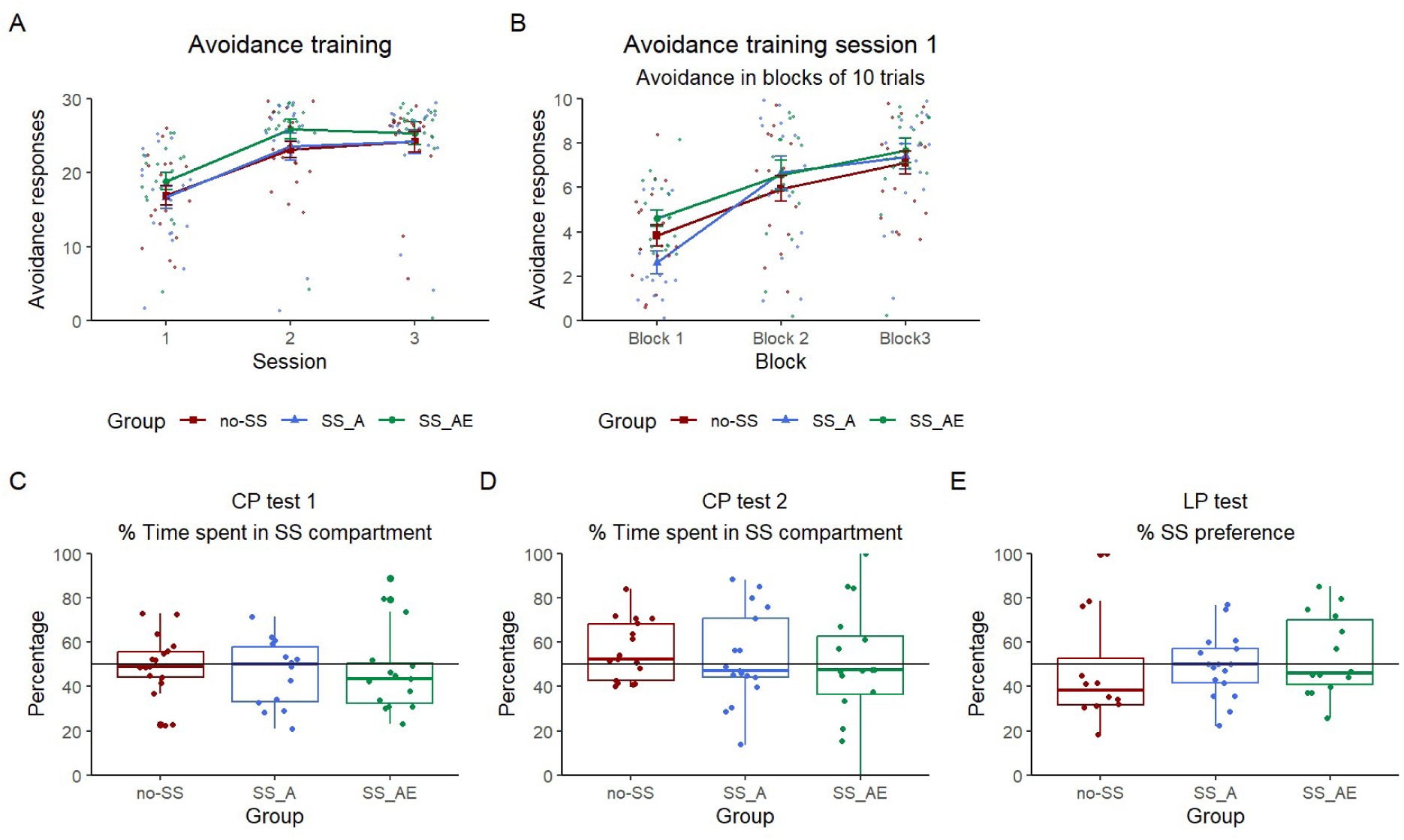
Experiment 5 avoidance training, compartment-preference and lever-preference tests. (A) Mean (± SEM) number of ARs for the SS_A (n = 18), SS_AE (n = 18) and no-SS (n = 18) groups. (B) Mean (± SEM) number of ARs of the first avoidance training session in blocks of 10 trials for all groups. (C) Boxplot of the percentage time spent in the SS-compartment of CP test 1 for rats of all groups. (D) Boxplot of the percentage time spent in the SS-compartment of CP test 2 for all groups. (E) Boxplot of the percentage SS preference of the LP test for all groups.

During CP test 1, as expected, rats of the no-SS group did not spend more time with the SS, compared to the CC (*t*(13) = 1.470, *p* = .165, *d* = 0.393; Figure 4C). In contrast with our hypothesis, rats of the SS_A group did not show a significant SS preference effect either (*t*(13) = −0.891, *p* = .805, *d* = −0.238). Similarly, during CP test 2, rats of both the no-SS group (*t*(13) = 1.725, *p* = .108, *d* = 0.461) and the SS_A group (*t*(13) = −0.310, *p* = .619, *d* = −0.083) did not show preference for the safety signal (Figure 4D). Finally, in the lever-preference (LP) test, three subjects were excluded from the analyses due to a failure to press each lever at least once (post-hoc criteria). In contrast with our hypothesis, rats of the SS_A group did not show a significant preference for the SS lever, compared to the CC lever (*t*(12) = −0.757, *p* = .768, *d* = −0.210; Figure 4E). Unexpectedly, rats of the no-SS group preferred the CC lever over the SS lever (*t*(11) = −2.878, *p* = .015, *d* = −0.831), even though the cues and levers were fully counterbalanced (white light and yellow light, left lever and right lever).

### 3.5. Experiment 5

Experiment 4 could not replicate the SS-enhanced learning that was observed in Experiment 3. To further examine why we were unable replicate our findings, a new experiment was set up using the same experimental procedure as in Experiment 4, except that an SS_AE group was added to further disentangle the role of avoidance vs. avoidance and escape responses in assigning a rewarding value to the safety signal. The SS_A group (n = 18) received the safety signal contingent upon avoidance responses, the SS_AE group (n = 18) received the safety signal contingent upon avoidance and escape responses, and the no-SS group (n = 18) did not receive the safety signal contingent upon responses during avoidance training. We expected the largest avoidance acquisition acceleration and strongest rewarding value of the safety signal in the SS_AE group.

The results showed a significant effect of Session (*F*(1.64,83.71) = 73.292, *p* < .001, ƞ*_p_²* = 0.590), but no significant effect of Group (*F*(2,51) = 0.765, *p* = .470, ƞ*_p_²* = 0.029), nor a significant Group by Session interaction effect (*F*(3.28,83.71) = 0.299, *p* = .843, ƞ*_p_²* = 0.012; Figure 5A), in contrast with our hypothesis. Similarly, when restricting the analysis to the data of the first avoidance training session, there was a significant effect of Block (*F*(2,102) = 62.980, *p* < .001, ƞ*_p_²* = 0.553), but no effect of Group (*F*(2,51) = 0.780, *p* = .464, ƞ*_p_²* = 0.030) nor a significant Group by Block interaction (*F*(4,102) = 2.205, *p* = 0.074, ƞ*_p_²* = 0.08; Figure 5B).

Subjects that failed to shuttle during the CP tests, and therefore did not experience each cue at least once, were excluded from the analyses. As expected, rats of the no-SS group did not spend more time with the safety signal, compared to the control cue, during either CP test 1 (n = 17; *t*(16) = −0.144, *p* = .888, *d* = −0.035; Figure 5C) or CP test 2 (n = 17; *t*(16) = 1.867, *p* = .080, *d* = .453; Figure 5D). In contrast with our hypothesis, rats of the SS_A group did not show a SS-compartment preference either, be it during CP test 1 (n = 14; *t*(13) = −0.911, *p* = .811, *d* = −0.244; Figure 5C) or CP test 2 (n = 16; *t*(15) = 0.187, *p* = .427, *d* = 0.047; Figure 5D). Likewise, rats of the SS_AE group did not show a SS-compartment preference during either CP test 1 (n = 16; *t*(15) = 0.094, *p* = .463, *d* = 0.024; Figure 5C) or CP test 2 (n = 16; *t*(15) = −0.036, *p* = .514, *d* = −0.009; Figure 5D). For the LP test, subjects that did not press each lever at least once were excluded from the analysis. In line with the data of the CP tests, we did not observe any significant SS preference effect (no-SS group: *t*(10) = −1.368, *p* = .201, *d* = −0.413, n = 11; SS_A group: *t*(16) = −0.198, *p* = .577, *d* = −0.048, n = 17; SS_AE group: *t*(13) = 0.819, *p* = .214, *d* = 0.219, n = 14; Figure 5E).

### 3.6. Experiment 6

Experiment 5 showed no significant effects related to avoidance acquisition or to SS-compartment and SS-lever preference. This is in contrast with the evidence for accelerated avoidance acquisition we observed in Experiment 3 for rats that received a safety signal after avoidance responses (SS_A). Moreover, Experiment 2 suggested SS approach behavior for rats of the SS_A group, although a control group was lacking in this particular experiment. Interestingly, the studies that yielded significant effects (Experiment 2 and 3) were conducted in the setup from Coulbourn Instruments, whereas the studies that did not yield significant effects (Experiment 4 and 5) were conducted in a Med Associates setup. To evaluate whether the effect of the safety signal is setup-dependent (setups may entail differences in CS and US properties, and in the sensitivity of behavior detection), Experiment 6 aimed to replicate Experiment 5 in the Coulbourn instruments setup, with the exception that the LP test was removed.

The results showed a significant effect of Session (*F*(1.74,78.44) = 52.311, *p* < .001, ƞ*_p_²* = 0.538), but no significant effect of Group (*F*(2,45) = 1.765, *p* = .183, ƞ*²* = 0.073) nor a significant Group by Session interaction effect (*F*(3.49,78.44) = 0.385, *p* = .793, ƞ*_p_²* = 0.017; Figure 6A), suggesting that avoidance learning occurred similarly for all groups. When restricting the analyses to the data of the first avoidance training session, we similarly found an effect of Block (*F*(1.51,68.07) = 8.050, *p* = .002, ƞ*_p_²* = 0.152), but no significant effect of Group (*F*(2,45) = 1.466, *p* = .242, ƞ*_p_²* = 0.061), nor a Group by Block interaction effect (*F*(3.03,68.07) = 0.821, *p* = .487, ƞ*_p_²* = 0.035; Figure 6B). Interestingly, visual exploration of the data pointed towards a trend of accelerated avoidance learning for rats of the SS_AE group, compared to the no-SS group. Therefore, an exploratory mixed ANOVA with repeated-measures factor Session and between-subjects factor Group (SS_AE and no-SS) was conducted. Despite the trend towards more avoidance responses for rats of the SS_AE group, the effect of Group remained non-significant (*F*(1,30) = 3.747, *p* = .062, ƞ*_p_²* = 0.111). As such, we were unable to replicate the results of Experiment 3, where the safety signal presented after avoidance responses was successful in accelerating avoidance learning. One potential explanation as to why we were unable to replicate the results of Experiment 3 could lie in the different safety signal stimuli that were used, although exploratory analyses (see Supplementary Materials) argue against this.

**Figure 6.**
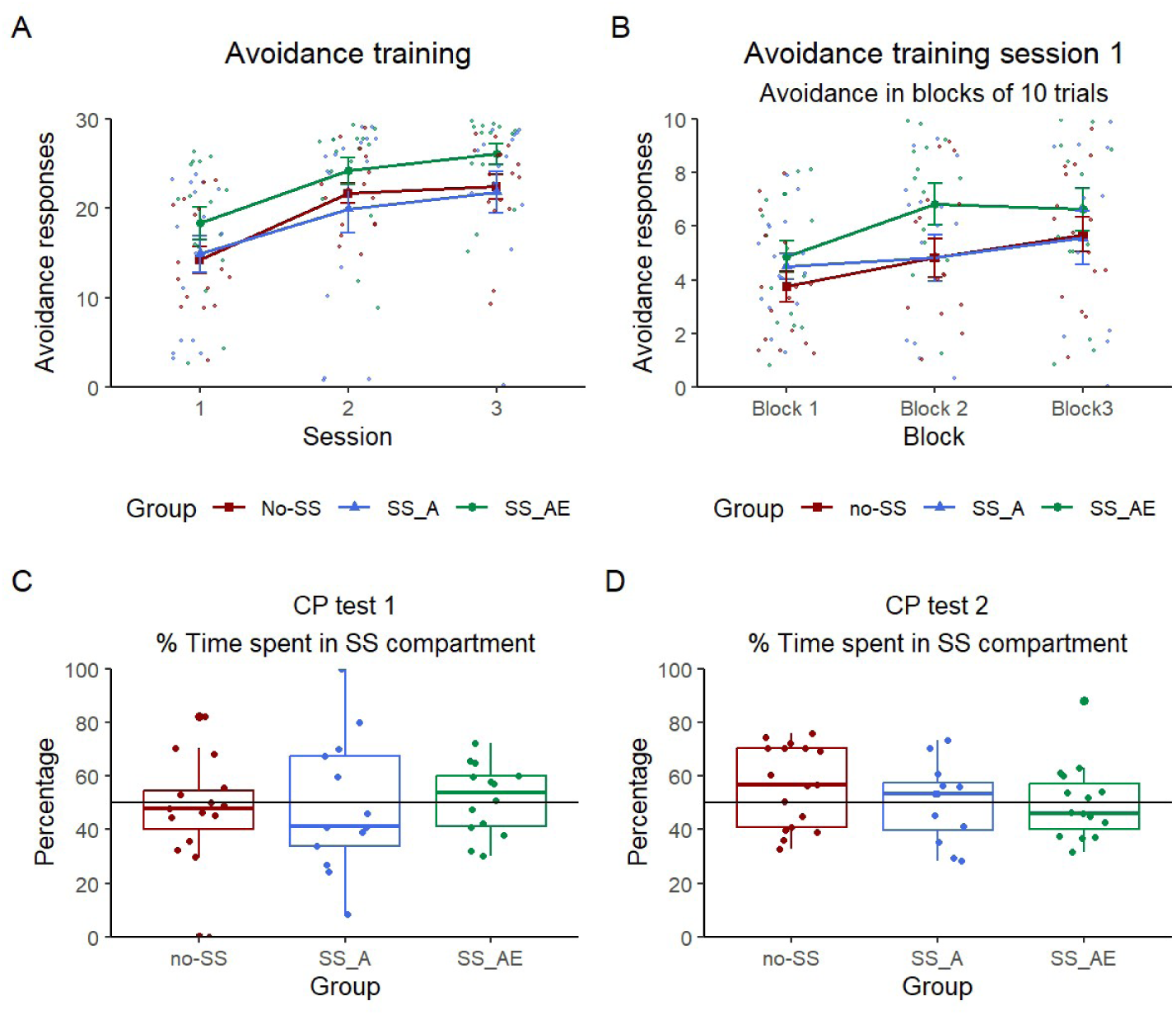
Experiment 6 avoidance training and compartment-preference tests. (A) Mean (± SEM) number of ARs for the SS_A (n = 16), SS_AE (n = 16) and no-SS (n = 16) groups. (B) Mean (± SEM) number of ARs of the first avoidance training session in blocks of 10 trials for all groups. (C) Boxplot of the percentage time spent in the SS-compartment of CP test 1 for rats of all groups. (D) Boxplot of the percentage time spent in the SS-compartment of CP test 2 for all groups.

Subjects that failed to shuttle during the CP tests, and therefore did not experience each cue at least once, were excluded from the analyses. During CP test 1, none of the groups showed a significant SS preference effect (no-SS group: *t*(14) = −0.524, *p* = .609, *d* = −0.135, n = 15; SS_A group: *t*(12) = −0.129, *p* = .550, *d* = −0.036, n = 13; SS_AE group: *t*(13) = 0.387, *p* = .352, *d* = 0.103, n = 14; Figure 6C). The same pattern of results was found in CP test 2 (no-SS group: *t*(15) = 1.471, *p* = .162, *d* = 0.368, n = 16; *t*(11) = 0.075, *p* = .471, *d* = 0.022, n = 12; SS_AE group: *t*(15) = 0.442, *p* = .332, *d* = 0.111, n = 16; Figure 6D).

## 4. Discussion

The present study aimed to assess whether the omission of an aversive event is inherently rewarding and thereby serves as a positive reinforcer for avoidance behavior (Cain, 2019). To this end, we adopted a behavioral approach to examine whether safety signals that coincide with avoidance-induced omissions of aversive events acquire rewarding properties. Previous research observed increased dopamine release in the nucleus accumbens during safety signals that signal successful avoidance of, or escape from, an aversive event (Oleson et al., 2012), in a manner similar to dopamine responses elicited by reward-predictive cues (Schultz, 2016). These findings raise the possibility that the brain processes the omission of an aversive event in a manner analogous to receiving a reward. However, given that dopamine is also implicated in processes unrelated to reward, such as salience and attentional modulation (Osorio-Gómez et al., 2022; Matzel & Sauce, 2023) and potentially also punishment (Amin et al., 2025), increased dopamine release alone cannot conclusively demonstrate that US omissions are processed as rewards. To address this limitation, we propose evaluating the behavioral responses elicited by cues associated with US omissions (i.e., safety signals) as a means to infer their potential rewarding properties.

Across six experiments, we investigated whether a safety signal (1) accelerates the acquisition of two-way active avoidance, (2) elicits approach behavior, and (3) motivates the acquisition of a novel instrumental response. Collectively, our findings indicate that safety signals can accelerate avoidance learning, but, as this effect was not consistently observed across all experiments, the boundary conditions under which this effect occurs remain to be clarified. Moreover, we were unable to observe significant effects on the compartment-preference task or on the lever-preference task across the majority of experiments, thereby questioning whether avoidance-contingent safety signals elicit reward-like behavior.

Experiments 1, 3, 4, 5 and 6 evaluated whether a safety signal can accelerate avoidance learning, in line with what has previously been observed using other avoidance tasks (Cándido et al., 1991; Dillow et al., 1972). We did observe more avoidance in rats that received a safety signal in Experiment 3 and Supplemental Experiment A, but not (or not significantly) in the other experiments. These findings suggest that safety signals can accelerate avoidance acquisition under some conditions, but the exact boundary conditions remain unclear. There were several small changes in experimental parameters across the different experiments that could account for the inconsistency of our findings, such as the number of training sessions (3 versus 5), the inclusion of a control cue, and the duration and intensity of the safety signal.

The inconsistency to replicate SS-enhanced avoidance learning is in contrast to previous studies that observed accelerated avoidance learning when a safety signal was included (Cándido et al., 1991; Dillow et al., 1972). An important difference between the studies described above and previous literature is the type of avoidance task that was used. The 2WAA task takes advantage of build-in species-specific defensive reactions (i.e., flight; Bouton and Fanselow, 1997), and is easier to acquire than avoidance tasks that involve the performance of a novel instrumental response in order to successfully avoid foot shock (Cain, 2019). Lever-press avoidance, for instance, requires shaping procedures and extensive training and, even then, performance will be lower compared to shuttle avoidance. Consequently, safety signals may be more relevant in contexts that require more complex behaviors. In our experiments, rats acquired the avoidance response readily, with asymptotic performance reached as early as the second or third training session. This rate of acquisition appears relatively fast compared to prior studies. For instance, Galatzer-Levy et al. (2014) identified four distinct behavioral trajectories that emerge during 2WAA learning. They differentiate between rapid avoiders who acquired the avoidance response on the first day (22%), modal avoiders, who reached criterion after three days (37%), slow avoiders, requiring five days of training (22%), and non-avoiders, who failed to acquire the response altogether despite extensive training (20%). In contrast, 79% of our animals reached asymptotic performance (defined as >80% avoidance) within three training days, which led us to shorten the training paradigm from five to three sessions in subsequent experiments. Furthermore, only 10 out of 246 animals (approximately 4%) failed to acquire the avoidance response entirely.

One potential explanation for the difference in learning speed between our study and that of Galatzer-Levy et al. (2014) may be the absence of an initial Pavlovian trial in our design. In our experiments, animals did not uniformly experience the CS–US pairing at the beginning of the task. In some cases, rats successfully performed avoidance responses during the first few CS presentations, thereby preventing any exposure to the US during early trials. This early success may have contributed to the perception that the aversive stimulus was avoidable, potentially enhancing motivation and facilitating rapid learning.

Experiments 2 through 6 investigated whether a safety signal elicits approach behavior following the acquisition of avoidance, as has previously been demonstrated for safety signals established via conditioned inhibition (Rogan et al., 2005) and for cues predicting natural rewards (Berridge & Robinson, 2003). A significant preference for the compartment associated with the safety signal was observed in Experiments 2 and 3, but not in Experiments 4, 5, and 6. Notably, the experiments in which a significant preference effect was observed lacked an appropriate control group, limiting our ability to determine whether the effect reflected a true preference for the safety signal. In Experiment 2, all animals were exposed to the safety signal during avoidance training, and in Experiment 3, even the control group displayed a preference for the safety signal, possibly driven by its novelty rather than its putative safety-related properties. Our findings stand in contrast to those observed for safety signals established through conditioned inhibition, as reported by Rogan et al. (2005). A key distinction, besides the differing paradigms used to establish the safety signal (avoidance learning vs. conditioned inhibition), lies in the nature of the safety signal itself. While Rogan et al. (2005), and similarly Fernando et al. (2014), used an auditory cue (a tone) as the safety signal, our experiments employed a visual stimulus (a light). Nonetheless, the use of light as a safety signal in avoidance learning is well-documented (Cándido et al., 1991; Feldman, 1977; Brennan et al., 2003), and prior research has demonstrated that light can function as a conditioned inhibitor, meeting both summation and retardation criteria following avoidance training (Cándido et al., 2004). Another critical difference between our study and those using conditioned inhibition, such as Rogan et al. (2005), concerns the predictive nature of the safety signal. In conditioned inhibition procedures, the safety signal reliably predicts the omission of the US, with a fixed temporal relationship between the cue and the absence of the US. In contrast, in avoidance paradigms, the safety signal is contingent on the animal’s behavior and coincides with the omission of the US. The duration of this omission depends on when the animal transitions to the safe compartment, which can vary across trials (up to the 20-second duration of the CS). Thus, rather than functioning as a classical predictor, the safety signal in avoidance learning serves more as a feedback cue that signals successful avoidance, rather than proactively signaling safety. The inability of our safety signal to elicit approach behavior was further supported by its failure to promote the acquisition of a novel instrumental lever-press response. This finding aligns with the results reported by Fernando et al. (2013), who similarly observed that a safety signal established via conditioned inhibition did not promote instrumental responding, in contrast to an appetitively trained conditioned stimulus. Collectively, these results suggest that safety signals derived from avoidance learning may not evoke reward-like behavioral effects, unlike safety signals established through conditioned inhibition, which have been shown to elicit approach behavior (Rogan et al., 2005).

In summary, our findings suggest that safety signals can accelerate avoidance learning under some conditions, but the circumstances under which this holds true remain unclear. Moreover, this also questions the robustness of the finding, as small changes to the experimental design, such as the duration of the safety signal or the inclusion of a control cue, may rendered the safety signal (in)effective. Finally, safety signals did not acquire rewarding properties that could be detected on a behavioral level, in contrast to appetitive conditioned cues or safety signals established by conditioned inhibition. Collectively, our findings argue against the notion that safety signals acquire rewarding properties by omissions of the aversive event.

## Authorship contribution

Laura Vercammen: conceptualization, data curation, formal analysis, funding acquisition, investigation, methodology, visualization, writhing – original draft, writing – review and editing; Tom Beckers: conceptualization, methodology, supervision, writing – review and editing; Bram Vervliet: conceptualization, methodology, supervision, writing – review and editing, funding acquisition; Laura Luyten: conceptualization, methodology, supervision, writing – review and editing

