## Supplementary Materials for "The rewarding properties of safety signals established by a two-way active avoidance task in male rats"

The Supplementary Material contains all supplemental tables (Supplementary Table 1 – 3) and preregistered analyses that were not included in the manuscript, as well as additional exploratory analyses.

### 1. Supplementary Tables

Suppl. Table 1. Overview of the experimental parameters of the avoidance training sessions. SS_AE groups received the safety signal contingent upon avoidance and escape responses, SS_A groups received the safety signal only after successful avoidance responses, and no-SS groups did not receive any safety signal during avoidance training.

| Exp | Groups | Acclimation | SS type | SS duration | Presentation of SS | ITI |
|---|---|---|---|---|---|---|
| 1 | SS_AE (n = 12) | 5 minutes | House light | 5 seconds | Compartment where rat shuttles into | 60 – 180 s |
| | no-SS (n = 12) | | | | | |
| 2 | SS_AE (n = 28) | 5 minutes | House light | 5 seconds | Compartment where rat shuttles into | 30 – 90 s |
| 3 | SS_A (n = 14) | 5 minutes | House light | 5 seconds | Compartment where rat shuttles into | 30 – 90 s |
| | no-SS (n = 14) | | | | | |
| 4 | SS_A (n = 14) | 10 minutes (with cue habituation) | House light or Yellow light (counterbalanced) | 10 seconds | Both compartments simultaneously | 30 – 90 s |
| | no-SS (n = 14) | | | | | |
| 5 | SS_AE (n = 18) | 10 minutes (with cue habituation) | House light or Yellow light (counterbalanced) | 10 seconds | Both compartments simultaneously | 30 – 90 s |
| | SS_A (n = 18) | | | | | |
| | no-SS (n = 18) | | | | | |
| 6 | SS_AE (n = 16) | 10 minutes (with cue habituation) | House light or Yellow light (counterbalanced) | 10 seconds | Both compartments Simultaneously | 30 – 90 s |
| | SS_A (n = 16) | | | | | |
| | no-SS (n = 16) | | | | | |
| Suppl Exp A | SS_A (n = 18) | 5 minutes | House light | 5 seconds | Compartment where the rat shuttles into | 30 – 90 s |
| | no-SS (n = 18) | | | | | |

Suppl. Table 2. Overview of the parameters used during the compartment-preference (CP) tests.

| General | | CP test 1 | | | CP test 2 | | |
|---|---|---|---|---|---|---|---|
| **Exp** | **Setup** | **Habituation** | **SS compartment assignment** | **Other compartment** | **Habituation** | **SS-compartment** | **Other compartment** |
| 2 | Modified shuttle box | 5 minutes | Position of rat after 5 minutes | No stimulus | 5 seconds | Same as CP test 1 | No stimulus |
| 3 | Modified shuttle box | 5 minutes | Counterbalanced | No stimulus | 5 seconds | Opposite to CP test 1 | No stimulus |
| 4 | Original shuttle box | 5 seconds | Counterbalanced | Control cue | 5 seconds | Opposite to CP test 1 | Control cue |
| 5 | Original shuttle box | 5 minutes | Counterbalanced | Control cue | 5 seconds | Opposite to CP test 1 | Control cue |
| 6 | Original shuttle box | 5 minutes | Counterbalanced | Control cue | 5 seconds | Opposite to CP test 1 | Control cue |
| Suppl Exp A | Modified shuttle box | 5 minutes | Counterbalanced | No stimulus | 5 seconds | Opposite to CP test 1 | No stimulus |

Suppl. Table 3. The presentation schedule of the cues (CC = control cue, SS = safety signal) during the 10-min habituation phase of the avoidance training sessions of Experiments 4, 5 and 6. Each cue was presented for 10 seconds.

| | 60 s | 85 s | 115 s | 155 s | 185 s | 220 s | 260 s | 290 s | 325 s | 365 s |
|---|---|---|---|---|---|---|---|---|---|---|
| SS | CC | / | CC | / | / | CC | / | CC | / | CC |
| no-SS | CC | SS | CC | SS | SS | CC | SS | CC | SS | CC |

### 2. Avoidance latencies

The average latency to avoid the aversive stimulus (US) was calculated for each avoidance training session per group and analyzed using mixed design ANOVAs with repeated-measures factor Session and between-subjects factor Group.

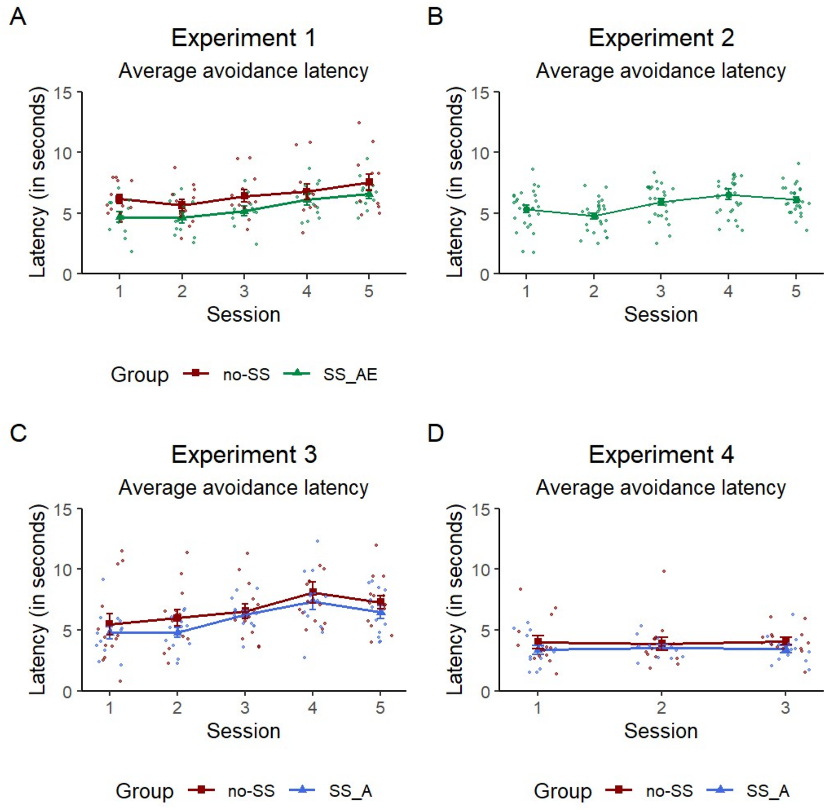


Supplemental Figure 1. The mean (± SEM) latency to avoid the aversive US for (A) Experiment 1, (B) Experiment 2, (C) Experiment 3, and (D) Experiment 4.

**Experiment 1.** We observed a significant effect of Session ($F(4,88) = 11.701$, $p < .001$, $\eta^2_p$ = 0.347) and a non-significant effect of Group ($F(1,22) = 3.880$, $p = .062$, $\eta^2_p$ = 0.150), with a numerical trend towards higher latencies for rats of the no-SS group (Suppl. Fig. 1A). There was no significant Group by Session interaction effect ($F(4,88) = 0.454$, $p = .769$, $\eta^2_p$ = 0.020).

**Experiment 2.** We observed a significant effect of Session ($F(2.71,70.54) = 6.183$, $p = .001$, $\eta^2_p$ = 0.192), suggesting that the latency to make an avoidance response increased with repeated training (Suppl. Fig. 1B).

**Experiment 3.** We observed a significant main effect of Session ($F(2.74,68.6) = 10.676$, $p <$ .001, $\eta^2_p$ = 0.299), but no main effect of Group ($F(1,25) = 1.173$, $p = .289$, $\eta^2_p$ = 0.045), nor a significant Group by Session interaction effect ($F(2.74,68.6) = 0.255$, $p = 0.841$, $\eta^2_p$ = 0.010; Suppl. Fig. 1C).

**Experiment 4.** We observed no significant main effect of Session ($F(2,52) = 0.034$, $p = .967$, $\eta^2_p = 0.001$), but no significant main effect of Group ($F(1,26) = 1.848$, $p = .186$, $\eta^2_p = 0.066$), nor a significant Group by Session interaction effect ($F(2,52) = 0.141$, $p = .869$, $\eta^2_p = 0.005$; Suppl. Fig. 1D).

### 3. Freezing

To evaluate freezing, we averaged the number of seconds of freezing across all even CS presentations (CS2, CS4, CS6,… CS30) and CS1 (except during the first training session, given that rats had not yet experienced the CS-US association at that point) for each avoidance training session per group and applied a mixed design ANOVA with repeated-measures factor Session and between-subjects factor Group. For Experiment 1, freezing was scored during every fifth CS presentation (CS1, CS5,… CS30), instead of during every even CS presentation (Moscarello & Ledoux, 2013). Although freezing is often represented as a percentage of CS duration in classical conditioning studies, we analyzed freezing in number of seconds because of the varying CS durations, in line with previous two-way active avoidance studies (e.g., Moscarello & Ledoux, 2013).

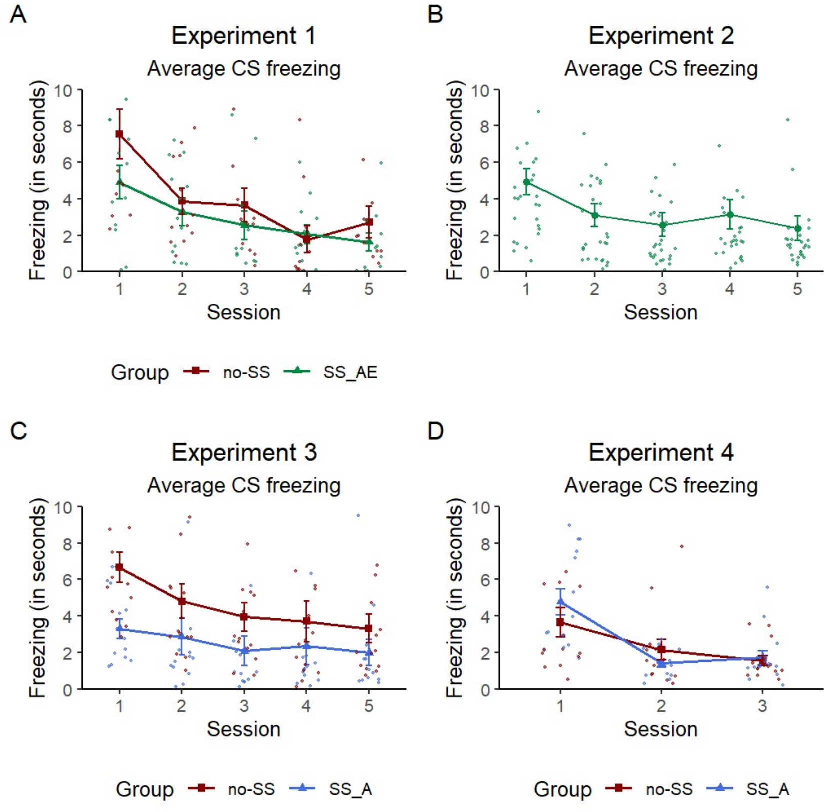


Supplemental Figure 2. The mean (± SEM) number of seconds of freezing for (A) Experiment 1, (B) Experiment 2, (C) Experiment 3, and (D) Experiment 4.

**Experiment 1.** We observed a significant main effect of Session ($F(2.92,64.31) = 18.922$, $p < .001$, $\eta^2_p = 0.462$), but no effect of Group ($F(1,22) = 1.180$, $p = .289$, $\eta^2_p = 0.051$), nor a significant Group by Session interaction ($F(2.92,64.31) = 1.793$, $p = .159$, $\eta^2_p = 0.075$; Suppl. Fig. 2A)

**Experiment 2.** We observed a significant effect of Session ($F(1.75,47.33) = 10.676$, $p < .001$, $\eta^2_p = 0.283$), suggesting that freezing decreased across training (Suppl. Fig. 2B).

**Experiment 3.** We observed a significant effect of Session ($F(2.4,59.88) = 7.133$, $p < .001$, $\eta^2_p = 0.222$), but no significant effect of Group ($F(1,25) = 3.448$, $p = 0.075$, $\eta^2_p = 0.121$), nor a significant Group by Session interaction effect ($F(2.4,59.88) = 1.406$, $p = 0.253$, $\eta^2_p = 0.053$), suggesting that freezing occurred similarly for both groups (Suppl. Fig. 2C).

**Experiment 4.** We observed a significant effect of Session ($F(1.48,38.58) = 26.375$, $p < .001$, $\eta^2_p = 0.504$), but no significant effect of Group ($F(1,26) = 0.108$, $p = 0.745$, $\eta^2_p = 0.004$) nor a significant Group by Session interaction effect ($F(1.48,38.58) = 2.710$, $p = .093$, $\eta^2_p = 0.094$), suggesting that there were no group differences in freezing (Suppl. Fig. 2D).

### 4. Additional preregistered analyses Experiment 2

A preregistered analysis was performed on the % time spent in the SS-compartment for each individual minute, for both CP test 1 and CP test 2. Time spent in the SS-compartment was significantly above 50 % for Minute 1 ($t(27) = 2.951$, $p = .003$, $d = 0.558$), Minute 3 ($t(27) = 2.129$, $p = .021$, $d = 0.402$) and Minute 5 ($t(27) = 2.394$, $p = .012$, $d = 0.453$), but not for Minute 2 ($t(27) = 1.325$, $p = 0.098$, $d = 0.250$) and 4 ($t(27) = 0.480$, $p = .318$, $d = 0.091$) of CP test 1. For CP test 2, time spent in the SS-compartment was significantly higher than 50% for Minute 1 ($t(27) = 3.014$, $p = .003$, $d = 0.570$) and Minute 4 ($t(27) = 4.184$, $p < .001$, $d = 0.791$), but not for Minute 2 ($t(27) = 1.155$, $p = 0.129$, $d = 0.218$), Minute 3 ($t(27) = 0.264$, $p = .397$, $d = 0.050$) and Minute 5 ($t(27) = 0.907$, $p = 0.186$, $d = 0.171$).

### 5. Additional preregistered analyses Experiment 3

To further test whether avoidance learning is accelerated when a safety signal is included, we conducted a preregistered exploratory analysis for which the avoidance responses were analyzed on a trial-by-trial basis by coding the data of the first avoidance training session in binary format (avoidance = 1, escape/escape failure = 0) and applying a generalized linear mixed-effects model to the binary outcome with fixed-effects parameters Trial, Group and the interaction between Trial and Group, and a random effects parameter Subject to account for within-subject variability due to the repeated measures design, using the R-package lme4 (https://cran.r-project.org/package=lme4). The logistic regression coefficients indicated a significant effect of Trial ($\beta = 0.035$, $SE = 0.013$, $Z = 2.709$, $p = .007$), suggesting that avoidance increased across trials. Moreover, there was a significant interaction between Trial and Group ($\beta$ = 0.051, *SE* = 0.020, *Z* = 2.550, *p* = .011), in line with the hypothesis that avoidance learning is enhanced when a safety signal is included.

A preregistered analysis was performed on the % time spent in the SS-compartment for each individual minute for both CP test 1 and CP test 2. For each individual minute, time spent in the SS-compartment was not significantly higher than 50 % for rats of the SS_A group (Minute 1: *t*(12) = 1.452, *p* = .086, *d* = 0.403; Minute 2: *t*(12) = 0.820, *p* = .214, *d* = 0.227; Minute 3: *t*(12) = -1.237, *p* = .880, *d* = -0.343; Minute 4: *t*(12) = 0.325, *p* = .375, *d* = 0.090; Minute 5: *t*(12) = 1.133, *p* = .140, *d* = 0.314). Time spent in the SS-compartment was not significantly different for each individual minute, except for Minute 1, for rats of the no-SS group (Minute 1: *t*(13) = 2.197, *p* = .047, *d* = 0.587; Minute 2: *t*(13) = 0.784, *p* = .447, *d* = 0.209; Minute 3: *t*(13) = 1.941, *p* = .074, *d* = 0.519; Minute 4: *t*(13) = 1.483, *p* = .162, *d* = 0.396; Minute 5: *t*(13) = 0.551, *p* = .591, *d* = 0.147). The fact that rats of the no-SS group spent more time in the SS-compartment during the first minute of the test might be due to the novelty of the cue. For CP test 2, rats of the SS_A group spent significantly more time in the SS-compartment, starting from Minute 3 onwards (Minute 1: *t*(12) = -3.602, *p* = .998, *d* = -0.999; Minute 2: *t*(12) = 1.483, *p* = .082, *d* = 0.411; Minute 3: *t*(12) = 1.780, *p* = .05, *d* = 0.494; Minute 4: *t*(12) = 2.069, *p* = .030, *d* = 0.574; Minute 5: *t*(12) = 1.898, *p* = .041, *d* = 0.527). For rats of the no-SS group, time spent in the SS-compartment was significantly different from 50 % for each Minute, except for Minute 3 (Minute 1: *t*(12) = -2.602, *p* = .023, *d* = -0.722; Minute 2: *t*(12) = 3.787, *p* = .003, *d* = 1.050; Minute 3: *t*(12) = 1.294, *p* = .220, *d* = 0.359; Minute 4: *t*(12) = 2.718, *p* = .019, *d* = 0.754; Minute 5: *t*(12) = 2.662, *p* = .021, *d* = 0.738).

**6. Additional preregistered analyses Experiment 4**

A preregistered exploratory generalized linear mixed model (GLMM) analysis was applied to the data of the first avoidance training session, to examine the effects of Trial and Group (reference group: no-SS group) on the binary avoidance data (avoidance = 1, escape/escape failure = 0), with a random intercept for each subject. The logistic regression coefficients indicated only a significant effect of Trial ($\beta$ = 0.057, *SE* = 0.013, *Z* = 4.436, $p < .001$), but no significant effects of Group, nor a significant interaction effect.

**7. Additional preregistered analyses Experiment 5**

A preregistered exploratory generalized linear mixed model (GLMM) analysis was applied to the data of the first avoidance training session, to examine the effects of Trial and Group (reference group: no-SS group) on the binary avoidance data (avoidance = 1, escape/escape failure = 0), with a random intercept for each subject. The logistic regression coefficients indicated only a significant effect of Trial ($\beta$ = 0.083, *SE* = 0.012, *Z* = 6.973, $p < .001$).

Preregistered analyses were conducted to evaluate whether rats of the SS_AE group and rats of the SS_A group show a higher SS preference effect during the CP tests, compared to rats of the no-SS group, and whether rats of the SS_AE group show a higher SS preference effect compared to rats of the SS_A group (SS_AE > SS_A > no-SS). For CP test 1, there was no significant difference between the SS_AE and the no-SS group ($t(31) = 0.157$, $p = .438$, $d = 0.055$), nor between the SS_A and the no-SS group ($t(29) = -0.611$, $p = .727$, $d = -0.221$), and the SS_AE and SS_A group ($t(28) = -0.589$, $p = .280$, $d = -0.215$). Similarly, we observed no group differences during CP test 2 (SS_AE – no-SS: $t(31) = -0.880$, $p = .807$, $d = -0.307$; SS_A – no-SS: $t(32) = -0.501$, $p = .690$, $d = -0.172$; SS_AE – SS_A: $t(31) = 0.405$, $p = .656$, $d = 0.141$).

Preregistered analyses were conducted to evaluate whether rats of the SS_AE group and rats of the SS_A group show a higher SS preference effect during the LP test, compared to rats of the no-SS group, and whether rats of the SS_AE group show a higher SS preference effect compared to rats of the SS_A group (SS_AE > SS_A > no-SS). We did not observe a significant difference between the SS_AE and the no-SS group ($t(24) = 0.828$, $p = .208$, $d = 0.326$). Similarly, we did not observe a significant difference between the SS_A and the no-SS group ($t(27) = 0.312$, $p = .379$, $d = 0.118$). Moreover, we expected that rats of the SS_AE group would show a higher SS preference effect, compared to rats of the SS_A group, since we expected that the safety signal would acquire more rewarding value when it is presented after both avoidance and escape responses. However, there was no significant difference between both groups ($t(29) = -0.799$, $p = .215$, $d = -0.288$).

**8. Additional preregistered analyses Experiment 6**

A preregistered exploratory generalized linear mixed model (GLMM) analysis was applied to the data of the first avoidance training session, to examine the effects of Trial and Group (reference group: no-SS group) on the binary avoidance data (avoidance = 1, escape/escape failure = 0), with a random intercept for each subject. The logistic regression coefficients indicated only a significant effect of Trial ($\beta = 0.042$, $SE = 0.012$, $Z = 3.556$, $p < .001$).

Preregistered analyses were conducted to evaluate whether rats of the SS_AE group and rats of the SS_A group show a higher SS preference effect during the CP tests, compared to rats of the no-SS group, and whether rats of the SS_AE group show a higher SS preference effect compared to rats of the SS_A group (SS_AE > SS_A > no-SS). There were no significant group differences for both CP test 1 (SS_AE – no-SS: $t(27) = -0.643$, $p = .263$, $d = -0.239$; SS_A – no-SS: $t(26) = -0.202$, $p = .421$, $d = -0.077$; SS_AE – SS_A: $t(25) = -0.296$, $p = .385$, $d = -0.114$) and CP test 2 (SS_AE – no-SS: $t(30) = 0.757$, $p = .773$, $d = 0.268$; SS_A – no-SS: $t(26) = 0.921$, $p = .817$, $d = 0.352$; SS_AE – SS_A: $t(26) = -0.231$, $p = .409$, $d = -0.088$).

**9. Supplementary analyses Experiment 6**

One potential alternative explanation as to why we were unable to replicate the results of Experiment 3 could lie in the different safety signal stimuli that were used, but exploratory analyses argue against this. In Experiment 3, all rats received the house light as a safety signal, whereas in Experiment 6, rats received either the house light or the yellow light as safety signal. To evaluate whether the type of stimulus could potentially influence the efficacy of the safety signal to accelerate avoidance learning, we performed an exploratory independent samples t-test. Specifically, the number of avoidance responses in the first avoidance training session was compared for the SS_A group of Experiment 3 (n = 14, all received the house light as safety signal) and the SS_A group that received the house light of Experiment 6 (n = 8). We observed a significant difference, with rats of Experiment 6 showing fewer avoidance responses during the first avoidance training session, compared to rats of Experiment 3, even though they both received the same cue as safety signal ($t(19) = 2.451$, $p = 0.024$, $d = 1.102$). Moreover, when exploratively comparing the SS_A group of Experiment 6 that received the house light safety signal (n = 8), with the SS_A group that received the yellow light safety signal (n = 8), we did not observe a significant difference in the number of avoidance responses performed during the first training session between both safety signal groups ($t(14) = -1.845$, $p = .086$, $d = -0.923$). In contrast, the findings even suggest a trend towards more avoidance for rats that received the yellow light as safety signal. Together, these findings suggest that the difference in cues that were used cannot explain why we were unable to replicate the results of Experiment 3.

### 10. Supplemental Experiment A

Supplemental Experiment A was conducted after Experiment 3, with the aim of testing an adapted behavioral protocol that was suitable for continuous *in vivo* microdialysis recordings. The experiment took place in the Coulbourn Instruments setup, as described in the Apparatus section of the Methods and Materials.

#### 10.1. Procedures

**Two-way active avoidance training.** Upon arrival in the lab, rats were randomly assigned to either the SS_A group (n = 18) or the no-SS group (n = 18). Two-way active avoidance training took place in the Coulbourn Instruments setup and proceeded as in Experiment 3, except that rats underwent three avoidance training sessions all on the same day, with a 20-min interval between each session. This change was added to the design in order to prepare for in vivo microdialysis. During the intervals, the rats were placed back in their home cage in the testing room.

**Compartment-preference test 1.** Twenty-four hours after the final avoidance training session, the rats were subjected to CP test 1. This test proceeded similarly as CP test 1 in Experiment 3.

**Compartment-preference test 2.** Twenty-four hours after CP test 1, the rats were subjected to CP test 2. This test proceeded similarly as CP test 2 in Experiment 3.

#### 10.2. Results

All rats were subjected to three avoidance training sessions that all took place on the same day, with a 20-minute interval between the sessions as described by Dombrowski et al., 2013. Rats were randomly assigned to either the SS_A group (n = 18), which received the safety signal after each successful avoidance response, or the no-SS group (n = 18), which did not receive the safety signal during avoidance acquisition.

We observed a significant main effect of Session, suggesting that the number of avoidance responses increased with repeated training ($F(1.56,51.45) = 44.968$, $p < .001$, $\eta_p^2 = 0.577$). Moreover, we observed a significant main effect of Group ($F(1,33) = 4.436$, $p = .043$, $\eta_p^2 = 0.118$), indicating that rats of the SS_A group showed more avoidance learning, compared to the no-SS group (Suppl. Fig. 3A), in line with the hypothesis that safety signals can accelerate avoidance learning.

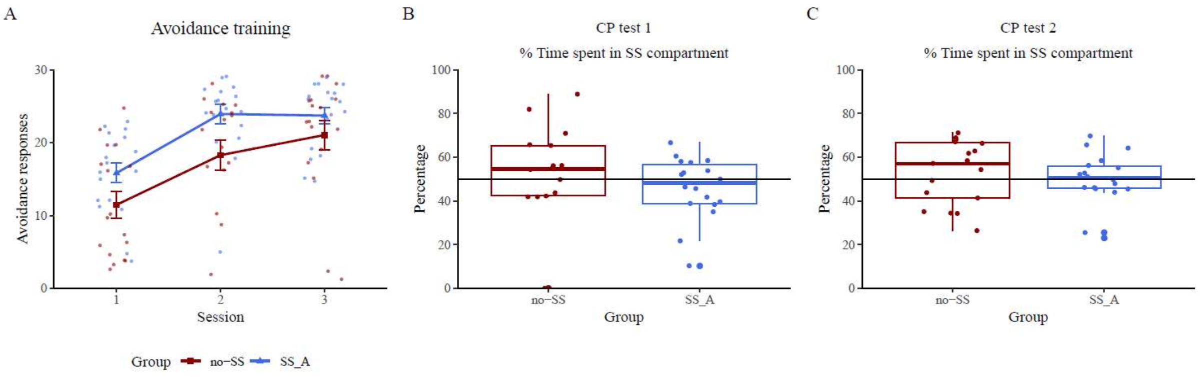


Supplemental Figure 3. Avoidance training and compartment-preference tests for Supplemental Experiment A. (A) Mean (± SEM) number of avoidance responses for the SS_A (n = 18) and the no-SS (n = 18) groups. (B) Boxplot of the percentage time spent in the SS-compartment of CP test 1 for both groups. (C) Boxplot of the percentage time spent in the SS-compartment of CP test 2 for both groups.

For the CP tests, neither group spent more time with the safety signal during CP test 1 (SS_A group: $t(17) = -1.183$, $p = .873$, $d = -0.279$; no-SS group: $t(16) = 0.195$, $p = .848$, $d = 0.047$; Suppl. Fig. 1B) or during CP test 2 (SS group: $t(17) = -0.006$, $p = .502$, $d = -0.001$; no-SS group: $t(16) = 0.863$, $p = .401$, $d = 0.209$; Suppl. Fig 1C). In sum, although the safety signal did enhance avoidance learning, it did not elicit approach behavior.

### 11. Pooled analyses

To further examine the effect of the safety signal on avoidance acquisition, we performed an exploratory pooled analysis over all experiments on the data of the first avoidance training sessions. A mixed ANOVA was applied with repeated-measures factor Block (Block 1, Block 2 and Block 3) and between-subjects factor Group (SS_AE, SS_A and no-SS). The data of each experiment was included, except for Experiment 2, which included only a single group of rats and was not set up to address potential group differences in avoidance acquisition.

We observed a significant effect of Block ($F(1.76,312.38) = 76.494$, $p < .001$, $\eta_p^2 = 0.301$), with the number of avoidance responses significantly increasing from Block 1 to Block 2 ($t_{\mathrm{Holm}} = -9.044$, $p < .001$) and from Block 2 to Block 3 ($t_{\mathrm{Holm}} = -2.785$, $p = .006$). Moreover, we observed a significant effect of Group ($F(2,178) = 4.553$, $p = .012$, $\eta_p^2 = 0.049$), with rats of the SS_AE group showing, on average, more avoidance responses than rats of the no-SS group ($t_{\mathrm{Tukey}} = 3.017$, p = .009). There was no significant difference in average avoidance responding for rats of the SS_A group and rats of the no-SS group ($t_{\mathrm{Tukey}} = 1.291$, $p = .402$). Together, these findings hint towards a general effect of the safety signal to enhance avoidance responding, when is presented contingent upon each avoidance and escape response, but not when it only follows successful avoidance responses.
